# H3K4me3 recruits the chromatin remodeling DILL-PICKLE-PEPPERCORN complexes to promoters in plants

**DOI:** 10.64898/2026.09.10.750567

**Authors:** Shuling Chen, Alberto Linares, Emilie Rannou, Nicole C. Härter, Chloé Champeyroux, Anja Schwarz, Elise Prellion, Alejandro Giraldo-Fonseca, Sibylle Pfammatter, Lixian Chen, Sisi Kang, Guohui Xie, Rui Liu, Jeff A. Long, Jiamu Du, Sylvain Bischof

**Affiliations:** Shenzhen Key Laboratory of Plant Genetic Engineering and Molecular Design, Institute of Plant and Food Science and Institute for Biological Electron Microscopy, Department of Biology, School of Life Sciences, Southern University of Science and Technology, Shenzhen, China; Department of Plant and Microbial Biology, University of Zurich, 8008 Zurich, Switzerland; Functional Genomics Center Zurich, University of Zurich/ETH Zurich, 8057 Zurich, Switzerland; Department of Biochemistry and Molecular Biology, International Cancer Center, Shenzhen University Medical School, Shenzhen 518060, China; State Key Laboratory of Genetics and Development of Complex Phenotypes, School of Life Sciences, Fudan University, Shanghai, China; Quzhou Fudan Institute, Quzhou, Zhejiang, China; Department of Molecular, Cell and Developmental Biology, University of California, Los Angeles, CA, USA

## Abstract

The plant CHD chromatin remodeler PICKLE (PKL) is a master regulator of cellular identity and differentiation, controlling developmental, hormonal and stress-response processes. Yet the molecular mechanisms underlying its function remain unclear. Here, we show that PKL forms three complexes, each composed of a protein of previously unknown function and one of three mutually exclusive novel DNAJ proteins that recruit HSP70-1. Simultaneous loss of all three DNAJs phenocopies the *pkl* mutant, indicating functional redundancy among PKL complexes. *In vitro* activity assays and cryo-electron microscopy reveal that PKL clamps nucleosomal DNA via its ATPase motor domain and recognizes H3K4me3 through its double chromodomain. Genome-wide profiling shows that H3K4me3 recognition positions PKL complexes at genic promoters to bidirectionally fine-tune gene expression. Together, these results provide structure-function insight into PKL recruitment and its control of developmental gene expression, and reveal a chaperone-coupled complex assembly that offers a broader perspective on how chaperone networks may support chromatin remodeling complexes across eukaryotes.

## Introduction

In eukaryotes, genomic DNA wraps around a histone octamer to form nucleosomes, which further pack into condensed chromatin^1^. DNA accessibility is controlled by chromatin remodelers, which use ATP hydrolysis to disrupt histone-DNA contacts, promoting more open or compact chromatin states and enabling downstream gene regulation^2,3^. Deficient chromatin remodeling is linked to developmental disorders, tumorigenesis, neurodegenerative and metabolic disease, and impaired plant growth and fertility^4–7^.

Chromatin remodelers fall into four families - switching defective/sucrose nonfermenting (SWI/SNF), imitation switch (ISWI), inositol requiring 80 (INO80), and chromodomain helicase DNA-binding (CHD)^2,3^ - that share a conserved SNF2-like ATPase domain but differ in accessory domains conferring distinct substrate recognition and regulatory properties. CHD remodelers feature N-terminal tandem chromodomain (CD) and a C-terminal composite DNA-binding domain^8,9^. The latter recognizes linker DNA flanking the nucleosome core particle (NCP), acting as a positional anchor that spaces nucleosomes evenly^10,11^, while the role of the double CD varies by subfamily. In subfamily I, human but not yeast CHD1 reads histone H3 lysine 4 trimethylation (H3K4me3)^12,13^ and is recruited with RNA polymerase II to active chromatin, promoting accessibility via nucleosome positioning or H3.3 deposition^14–16^. In subfamily II, the double CD are instead repurposed for direct DNA binding, while tandem N-terminal plant homeodomain (PHD) fingers confer H3K4me0/H3K9me3-binding specificity in CHD4^17^. CHD3/CHD4 form part of the Nucleosome Remodeling and Deacetylase (NuRD) complex, coupling remodeling with histone deacetylation to repress transcription^18–20^. The domain composition and functional diversity of CHD remodelers thus underlie their dual roles in transcriptional activation and repression, depending on molecular context and interacting partners.

The *Arabidopsis thaliana* genome encodes four CHD remodelers: the CHD1 homolog CHR5 and three CHD3 homologs, PICKLE (PKL), PICKLE RELATED 1 (PKR1), and PICKLE RELATED 2 (PKR2)^21,22^. Unlike PKR2, both PKL and PKR1 possess a PHD finger, reflecting architectural diversification within plant CHD subfamily II^21,22^. In rice, but not *Arabidopsis*, the recombinant CHD3 PHD finger and double CD bind H3K27me3 and H3K4me2 peptides, respectively^23,24^, and genetic and chromatin immunoprecipitation followed by sequencing (ChIP-seq) studies have established a complex relationship between PKL and these anti-correlated marks: PKL is enriched at repressed, H3K27me3-marked genes as well as active, H3K4me2/3-marked loci, and both modifications are redistributed in *pkl* mutants alongside the corresponding gene expression changes^23–28^. Consistent with this duality, loss of PKL reduces nucleosome density at H3K27me3 spreading regions^23^ but increases nucleosome occupancy genome-wide^29^. Reflecting these broad regulatory roles, *pkl* mutants display pleiotropic phenotypes, with PKL implicated in cellular identity and differentiation^25,26,30^, embryonic-to-vegetative transition^25,26,30,31^, heteroblasty^32^, flowering time^33^, root meristem activity^25,26^, lateral root, seed and endosperm formation^27,34,35^, light and hormone signaling^30,36^, and abiotic stress responses^37,38^.

Despite PKL’s substantial impact on chromatin organization, how it assembles as a functional unit and is targeted to specific genomic loci has remained unresolved^39^. Here, we show that PKL does not act alone, but nucleates three distinct, mutually exclusive multi-subunit complexes, each comprising PKL, a previously uncharacterized protein we name PEPPERCORN (PEPE), and one of three partially redundant DNAJ proteins (hereafter DILL1-3) that recruit the chaperone HSP70-1. Simultaneous loss of all three DILLs phenocopies *pkl*, demonstrating that the three complexes act redundantly, and our data indicate that PKL itself serves as the scaffold that independently recruits PEPE and each DILL. Our *in vitro* binding assay identified that PKL uses its double CD to bind to H3K4me3. Using cryo-EM, we resolved the structure of PKL bound to a H3K4me3-modified nucleosome core particle, showing that the PKL ATPase domain clamps nucleosomal DNA at the canonical superhelix location 2 (SHL2) position for chromatin remodeling while its double CD directly reads H3K4me3 - a binding mode we further validated biochemically and by structure-guided mutagenesis *in vivo*. Together, these results provide structure-function insight into PKL recruitment and PKL-mediated control of developmental gene expression, and reveal a chaperone-coupled complex assembly that offers a new perspective on how chaperone networks may more broadly support chromatin-remodeling complexes across eukaryotes.

## Results

### PKL interacts with a protein of unknown function and DNAJs

To investigate PKL’s mechanism of action, we expressed *pPKL::gPKL::2xYFP-3xFLAG-3’utrPKL* (PKL-YF) in the *pkl-10* background (referred as *pkl*)^27,40,41^. PKL-YF fully complemented the leaf-shape and delayed juvenile-to-adult transition phenotypes of *pkl* (Supplementary Fig. 1a,b) and localized constitutively to the nucleus across all examined cells and tissues (Fig. 1a and Supplementary Fig. 2), consistent with the ePlant expression database^42^. Immunoprecipitation followed by mass spectrometry (IP-MS) of PKL-YF in flowers identified a protein of unknown function, hereafter PEPPERCORN (PEPE), and three uncharacterized J-domain proteins, DILL1-3, as interactors (Fig. 1b,c and Supplementary Data 1,2). These interactions were resistant to benzonase treatment^43^, indicating that PKL complexes assemble independently of DNA.

**Fig. 1.**
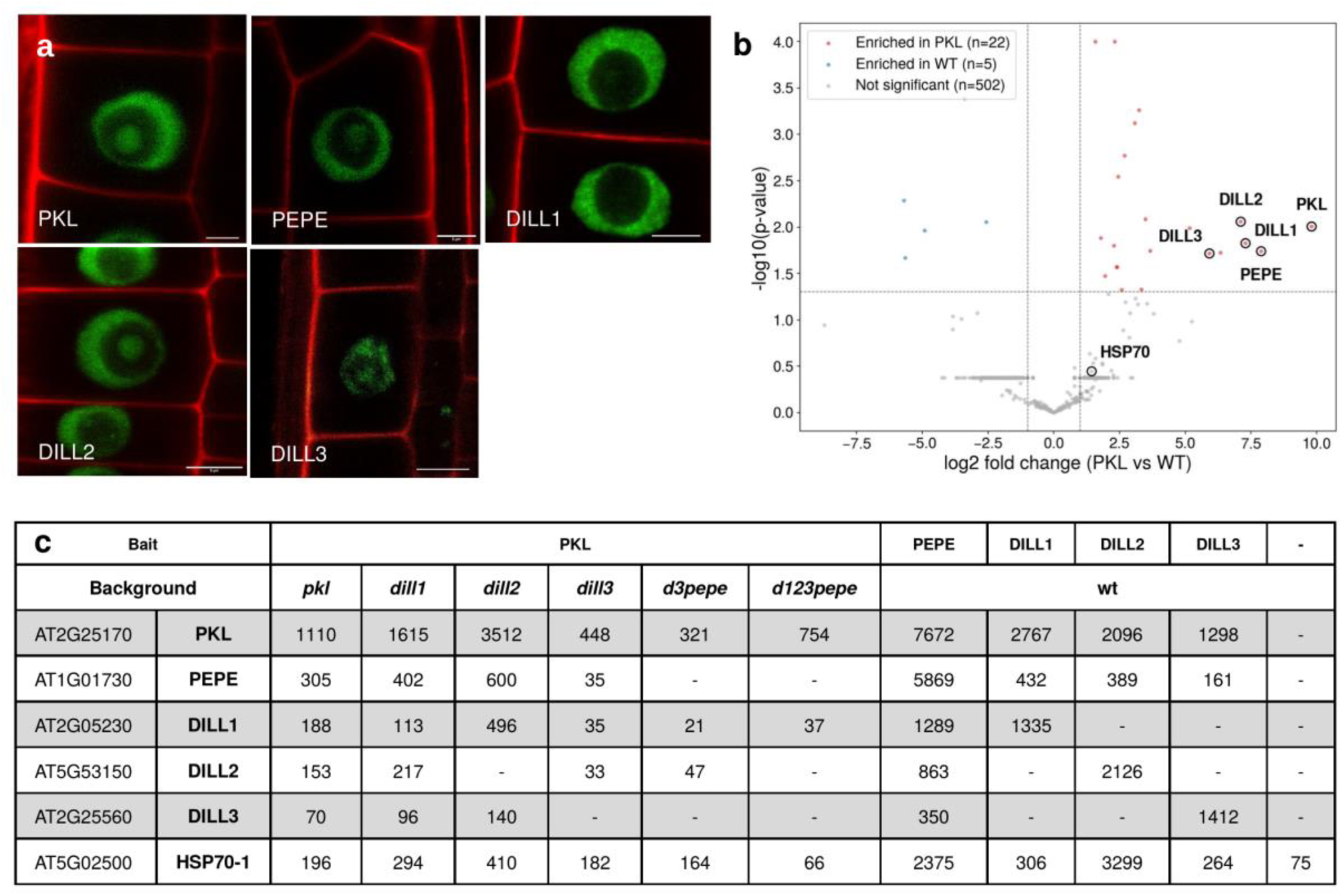
PKL forms three mutually exclusive complexes. a, Confocal microscopy showing the localization of 2xYFP-3xFLAG (YF) tagged proteins driven by their respective endogenous promoter in the root elongation zone (green). Cell walls were labelled with propidium iodide (red). Bar = 5 µm. **b,** Vulcano plot displaying the interaction partners of PKL-YF. Enrichment is significant if log2 fold-change| > 1 and P-value < 0.05. **c**, Table presenting the average spectral counts detected in two IP-MS replicates for each YF-tagged bait.

Reciprocal IP-MS on YF-tagged PEPE and DILL1-3, expressed under their endogenous promoters and localizing to the same nuclear compartments as PKL-YF (Fig. 1a and Supplementary Fig. 2), confirmed all three interactions with PKL (Fig. 1c). The DILLs did not co-immunoprecipitate with one another, indicating that PKL, PEPE, and each DILL assemble into three mutually exclusive complexes. PEPE encodes a single-copy, 224-residue protein with no homology to annotated domains. Structural prediction^44^ revealed a 55-residue, neutrally charged, leucine-rich helix-turn-helix (HTH) motif consistent with a protein-protein interaction or dimerization interface^45^ rather than a canonical DNA-binding HTH (Supplementary Fig. 3a,b). While PEPE is a plant-specific innovation, DILL1-3 instead belong to the HSP40/DNAJ family (Supplementary Fig. 3c,d), sharing only ∼36% amino acid identity but retaining canonical class A architecture - an N-terminal J-domain, G/F-rich linker, zinc finger-like region, and C-terminal substrate-binding domain. Among the 120 DNAJs encoded in the *Arabidopsis* genome, only five, including the three DILLs, carry a C-terminal DUF3444 of unknown function^46^. DILL1-3 form an independent phylogenetic clade (Supplementary Fig. 3e), distinct from DNAJs recently implicated in DNA methylation-reading complexes^47,48^, and no additional DNAJs were enriched in the PKL, PEPE, or DILL1-3 IP-MS datasets (Supplementary Data 1). These results identify PEPE and the DILLs as plant-specific regulatory components of CHD subfamily II.

### Redundant DILLs are required for PKL-mediated transcription

To assess the contributions of PEPE, DILL2 and DILL3 to plant development, we isolated T-DNA lines lacking full-length transcripts (Supplementary Fig. 3a,d,f,g). As the DILL1 locus contains two tandem duplications with identical gDNA and protein sequences, we used CRISPR-Cas9 to delete both DILL1 J-domains simultaneously. While *pepe* and *dill2* mutants were indistinguishable from wild type (WT), *dill1* and *dill3* displayed *pkl*-like leaf shapes and delayed flowering, implicating both in PKL function (Fig. 2). Higher-order *dill12*, *dill13*, *dill23* and *dill123* mutants, generated by cross-fertilization, showed progressively *pkl*-like phenotypes with increasing DILL loss, demonstrating genetic redundancy among *DILL1-3* (Fig. 2). Crossing *pepe* with single, double or triple *dill* mutants did not enhance any parental phenotype.

**Fig. 2.**
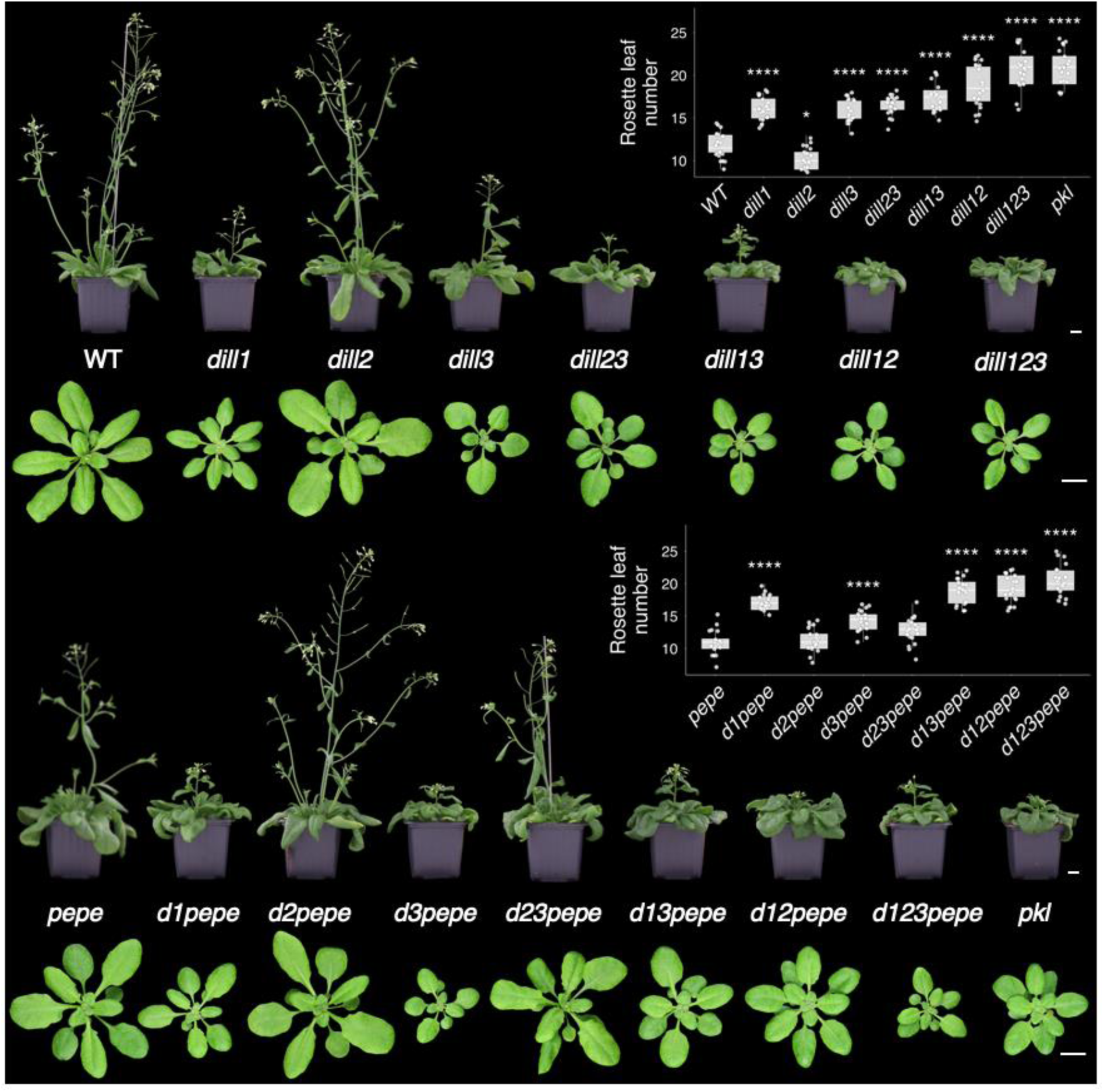
DILL1-3 are genetically redundant with PKL. Plant phenotype and quantification of the juvenile-to-adult transition by counting the number of rosette leaves at the time of bolting of WT, *pkl*, *dill1-3*, *pepe* and high-order mutant combinations. Bar = 1 cm. * and **** denote adjusted P-value < 0.05 and 0.0001.

RNA-seq of single and higher-order mutants (Fig. 3) showed that differentially expressed transcripts corresponded predominantly to protein-coding genes, with heterochromatic transposable elements remaining silent (Supplementary Data 3). The ratio of up- to down-regulated genes was similar across most backgrounds, indicating that DILL-PKL-PEPE complexes function in both transcriptional activation and repression (Fig. 3a). Although the DILLs could not fully compensate for one another, the number of differentially regulated genes increased progressively from single to double to triple *dill* mutants, supporting partial functional redundancy. Consistent with the phenotypic data (Fig. 2), the loss of PEPE neither strongly affected gene expression nor enhanced the transcriptional changes seen in *dill123*. PCA and clustering analyses showed that *dill1* and its combinations had the largest transcriptional impact among the *dill* mutants (Fig. 3b-d and Supplementary Fig. 4). Transcriptional changes in *dill123* and *dill123 pepe* remained milder than in *pkl*, suggesting that PKL retains partial functionality upon loss of its interaction partners, or that PKL has DILL- and PEPE-independent functions. Together, these genetic and genomic data establish reliable loss-of-function alleles and reveal partial redundancy among DILL1-3 within the PKL pathway.

**Fig. 3.**
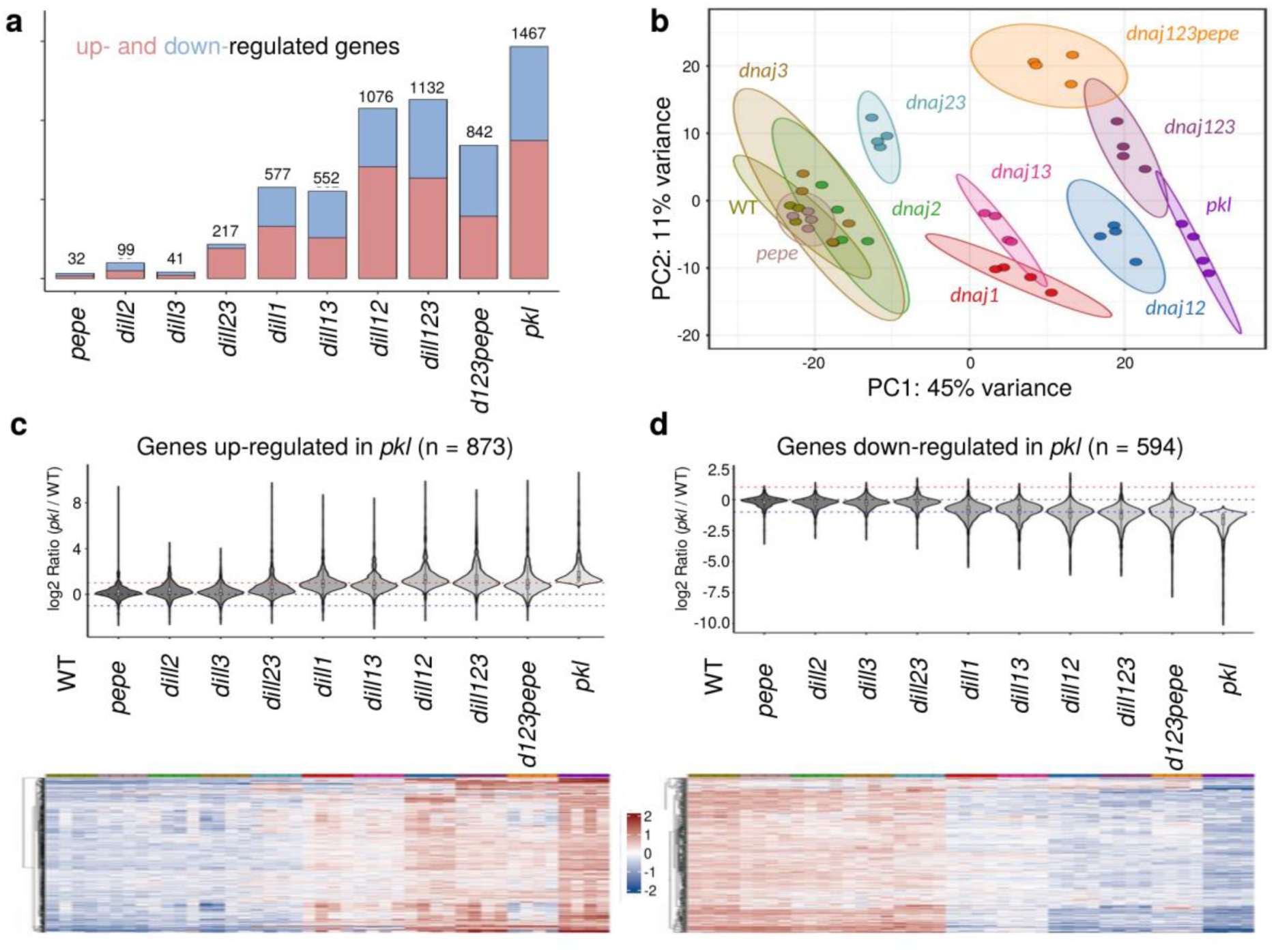
DILL1-3 are transcriptionally redundant with PKL. a, Bar graph of significantly up- and down-regulated genes in genotypes analyzed by RNA-seq (P < 0.05; fold-change > 2). **b,** PCA analysis of profiled transcriptomes. **c,d,** Violin plots and heatmaps showing gene expression in *pepe*, *dill1-3* and *pkl* mutant combinations for the genes differentially regulated in *pkl* (fold-change > 2 and P-value < 0.05).

### The DILL J-domain recruits HSP70-1 to PKL complexes

To resolve the complex hierarchy, we introgressed tagged PKL into *dill1*, *dill2*, and *dill3* single mutants, respectively, and performed IP-MS (Fig. 1c). The edited *dill1* line retained a truncated DILL1 protein lacking the N-terminus and J-domain (Supplementary Fig. 3g,h), yet this was sufficient for PKL interaction, indicating that the DILL1 C-terminus mediates PKL binding. In the *dill2* and *dill3* backgrounds, DILL2 and DILL3 were, respectively, not immunoprecipitated with PKL-YF, confirming these as likely null alleles. PEPE interacted with PKL-YF in all single *dill* backgrounds, showing that the DILLs are dispensable for this association. Likewise, PKL-YF pulldowns in a *dill3 pepe* double mutant still recovered DILL1 and DILL2, indicating that PEPE is not required for DILL2/3-PKL interactions. Together, these results show that PKL serves as the scaffold that independently recruits PEPE and the DILLs.

IP-MS identified HSP70-1 (AT5G02500) as significantly enriched with PKL, PEPE, and DILL1-3 relative to WT (fold-change = 10.9×, P = 0.0004; Fig. 1c and Supplementary Data 4). DNAJs act as HSP70 co-chaperones, binding client proteins and recruiting HSP70 via a conserved HPD tripeptide in their J-domain to stimulate its ATPase activity and mediate protein folding, disaggregation, or complex assembly/disassembly^46,49,50^. Neither loss of PEPE nor of individual DILLs reduced HSP70-1 enrichment, but in a *dill123 pepe* quadruple mutant, HSP70-1 abundance dropped to WT levels, confirming HSP70-1 as a bona fide component of the DILL-PKL-PEPE complexes. Together, IP-MS and genetics show that PKL independently anchors PEPE and the DILLs, and that the DILLs in turn recruit HSP70-1 to the complex.

### DILL-PKL-PEPE complexes are enriched at active promoters

The dispersed nuclear localization of DILL-PKL-PEPE complexes suggested binding to euchromatin (Fig. 1a). Indeed, ChIP-seq of tagged PKL, PEPE, and DILL1-3 showed enrichment on chromosomal arms and co-accumulation at genic promoters upstream of the transcriptional start site, a pattern typical of CHD remodelers^51^ (Fig. 4a-d and Supplementary Data 5), with depletion at centromeric and transposable-element regions, indicating that these complexes principally target protein-coding genes (Fig. 4a,b). PKL, PEPE, and DILL1-3 co-enriched at shared binding sites and target genes (Fig. 4e and Supplementary Fig. 5a,b), consistent with their physical association. PKL bound over 36% of all *Arabidopsis* genes, supporting a role as a widespread chromatin regulator (Fig. 4f). PKL-bound genes were highly expressed in WT, whereas unbound genes showed lower expression. Genes both bound by PKL and significantly up- or down-regulated in *pkl* were also highly expressed, confirming that PKL acts as both a transcriptional repressor and an activator at active loci. Metaplot analyses confirmed that PKL, PEPE, and DILL1-3 enrichment positively correlated with transcription (Fig. 4g and Supplementary Fig. 5c), and binning PKL-bound genes by expression level showed that PKL, PEPE, DILL1-3, and H3K4me3 - a hallmark of active transcription - all scaled positively with expression (Fig. 4h,i and Supplementary Fig. 5e). These results point to a recruitment mechanism for PKL guided by transcriptional activity or a specific epigenetic signature.

**Fig. 4.**
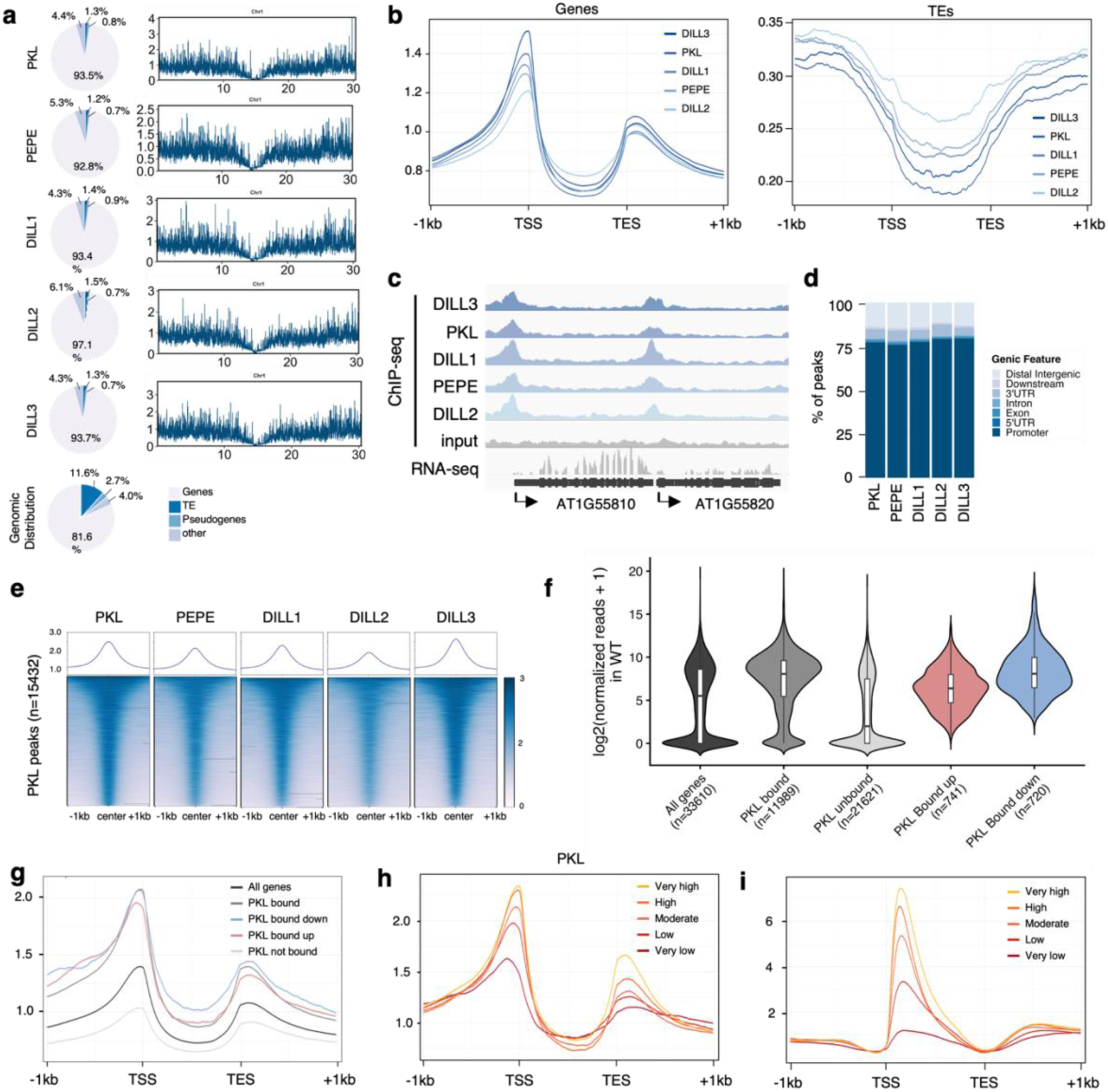
DILL-PKL-PEPE complex components co-localize at active promoters. a, Pie chart and chromosome view showing the distribution of mapped reads from PKL, PEPE and DILL1-3 ChIP-seq. **b,** Metaplot of PKL, PEPE and DILL1-3 over all protein-coding genes and transposable elements (TEs). **c,** IGV screenshot of ChIP-seq datasets and RNA-seq in WT. **d,** Stacked bar displaying the distribution of called peaks across genetic features. **e,** Metaplot and heatmap of PKL, PEPE and DILL1-3 enrichment at called PKL peaks. **f,** Violin plot presenting the expression in WT of all genes, genes bound or not by PKL, and PKL bound up- or down-regulated genes in *pkl*. **g,** Metaplot of PKL enrichment at the categories defined in f. **h,i,** Metaplot of PKL and H3K4me3 enrichment in five groups of active genes in WT defined according to their expression level.

### PKL binds to H3K4me3 by its double chromodomain

PKL was reported to associate with H3K4me2/3 and to regulate H3K27me3 deposition^23–29^, which prompted us to identify the potential histone modification recognized by PKL. Rice CHD3, the ortholog of *Arabidopsis* PKL, directly interacts with H3K4me2 and H3K27me3 via its CD and PHD finger, respectively^24^. The CD of human CHD1 was reported to directly bind to H3K4me3^12,13^, while the second PHD finger of human CHD4 directly recognizes the combined histone modification state of H3K4me0K9me3^17^. Similar to other CHD family chromatin remodelers, *Arabidopsis* PKL is a large multiple domain protein constituted by an N-terminal regulatory module of a plant PHD finger and two CDs, a central chromatin remodeling ATPase motor domain, and a C-terminal DNA-binding module of a Sant plus a Slide dual domains (Fig. 5a). Considering that the CD belongs to the canonical “Royal family” of histone methylation readers^52^, while the PHD finger constitutes a structurally distinct class of methyllysine-binding zinc-finger reader domains^53^, we tested their histone mark binding capacity by a Bio-layer interferometry–based *in vitro* screening assay. The various methylation states of common histone modification sites, including H3K4, H3K9, H3K27, and H3K36, were examined. The recombinant PKL double CD bound to the H3K4me3, H3K4me2, and H3K4me1 peptides with binding affinities of 7.7 µM, 17.7 µM, and 45.8 µM, respectively, but very weak or no detectable binding to other tested histone peptides, suggesting a binding preference for the higher methylation status of H3K4 site and consistent with the *in vivo* data that the PKL associates with H3K4me2/3 genome-wide^29^. Meanwhile, the PKL PHD finger showed very weak or no detectable binding to all the tested histone. Overall, our *in vitro* binding assay demonstrates that the double CD, but not the PHD finger, of PKL binds strongly to H3K4me3, suggesting a direct histone modification-based regulation of PKL.

**Fig. 5.**
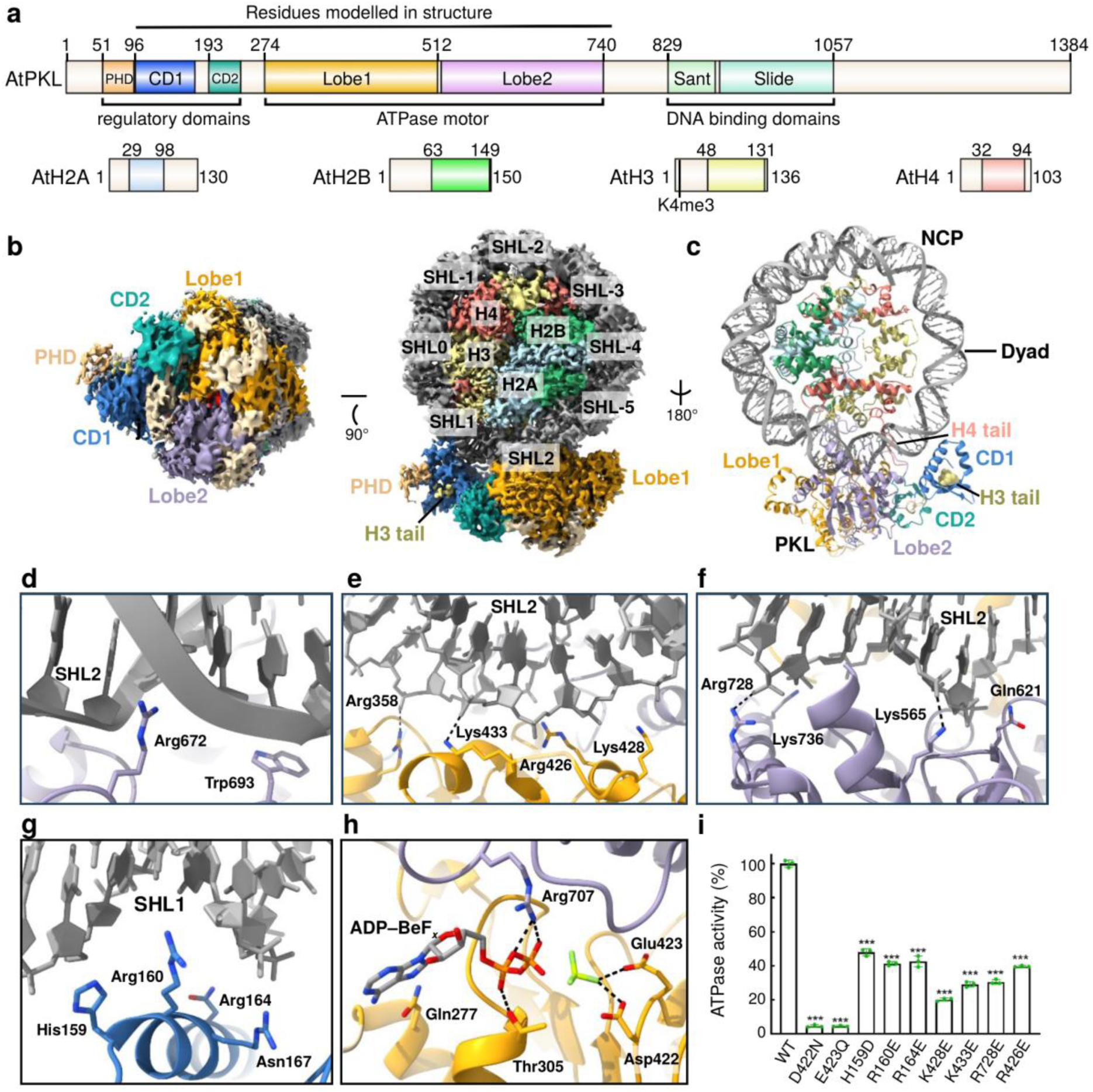
Structural insights into PKL and ATPase activity. **a,** Domain architecture of *A. thaliana* PKL and histone proteins used in this research. **b,** Two views of the cryo-EM map of the PKL-NCP complex related by 90°. **c,** Overall structure of PKL-NCP complex. **d-f,** Interaction details between the ATPase domain of PKL and nucleosomal DNA around SHL2 with the interacting residues and hydrogen bonds highlighted in stick and dashed-lines, respectively. **g,** Interaction details between the CD1 of PKL and nucleosomal DNA around SHL1. **h,** The active site conformation and ADP-BeF*x* binding. **i,** *In vitro* assays of representative PKL mutants affecting catalytic activity and DNA binding. The data are presented as means ± s.d. (*n* = 3 independent experiments); \*\*\**P* < 0.001 (two-tailed Student’s t-test). Individual data points corresponding to each mutant are shown as dots. The P-values for each mutant are as follows: D422N, 1.9 × 10-7; E423Q, 1.5 × 10-7; H159D, 6.0 × 10-6; R160E, 1.7 × 10-6; R164E, 1.3 × 10-5; K428E, 4.2 × 10-7; K433E, 1.1 × 10-6; R728E, 1.0 × 10-6; R426E, 9.8×10-7.

### Structure of PKL in complex with H3K4me3-modified NCP

To investigate the molecular basis of chromatin remodeling by PKL, we carried out structural and biochemical studies. The unmodified or H3K4me3-modified *Arabidopsis thaliana* NCPs were reconstituted using *Arabidopsis* histone proteins (Fig. 5a) and a 226-bp DNA consisting of 40-bp flanking DNA on both sides and a central 146-bp Widom 601-derived DNA as described before^54–57^. Consistent with a previous report^22^, the recombinantly expressed full-length PKL showed unambiguous activity in our ATPase activity-based chromatin remodeling assay (Supplementary Fig. 6a,b). The cryo-EM structure of PKL in complex with an H3K4me3-modified NCP and ADP-BeF_x_, a pre-hydrolytic-state ATP analog, was determined in two forms (Supplementary Fig. 6,7). The form one complex structure, consisting of one PKL bound to one NCP, was determined at a resolution of 3.14 Å, while the form two complex structure, consisting of two PKLs symmetrically bound on either side of one NCP, was determined at a resolution of 3.05 Å (Fig. 5b,c, Supplementary Fig. 6c). Similar to the *Arabidopsis* DDM1-NCP complex and other reported symmetric remodeler-NCP structures^55^, the two PKL-NCP binding interfaces of the 2:1 complex mirror that of the 1:1 complex (Supplementary Fig. 8). The local resolution of PKL was better in the 1:1 complex than in the 2:1 complex (Supplementary Fig. 6f,g), and since the biological significance of the 2:1 complex remains unclear, we focus on the 1:1 complex in the following discussion.

The PKL ATPase domain, consisting of the canonical Lobe1 and Lobe2, and CD1 are bridged by CD2 (Fig. 5b,c). The PHD finger has a map too weak to be modeled (Fig. 5b,c). However, the density could clearly locate the PHD finger to a position adjacent to the CD1, but the PHD finger has no direct contact with the CD2, the ATPase domain, and the NCP (Fig. 5b). We reconstituted the NCP with DNA possessing 40-bp flanking sequences on both the NCP entry and exit sides, which is long enough for the Sant and Slide DNA-binding domains to bind in other reported CHD-NCP structures^57^. However, we observed neither the flanking DNA sequences at the NCP entry and exit sides nor the Sant and Slide DNA-binding domains in our cryo-EM map (Fig. 5b), suggesting a flexible conformation of this subcomplex. Thus, these segments were not built. The H3K4me3-modified H3 tail bound to the CD1, with only H3T3 and H3K4me3 being traced and modeled.

### Recognition of NCP DNA by PKL

PKL ATPase domain Lobe1 and Lobe2 clamp at a canonical chromatin remodeler binding position of the nucleosomal DNA, the SHL2, with extensive interactions (Fig. 5b,c). A series of positively charged residues of both the Lobe1 and Lobe2 of the PKL ATPase domain approach the nucleosomal DNA backbone to form hydrogen bonding and salt bridge interactions (Fig. 5d-f). While CD2 and PHD finger do not interact with the NCP (Fig. 5b), a positively charged residues-enriched α-helix of PKL CD1, including His159, Arg160, Arg164, and Asn167, approaches the minor groove of the SHL1 DNA of the NCP to interact with the DNA backbone by hydrogen bonding and salt bridge interactions (Fig. 5g). The ADP-BeF_x_ is bound in the nucleotide binding pocket of the PKL ATPase domain with a clear map (Supplementary Fig. 6f). The Lobe1 and Lobe2 extensively interact with the ADP-BeFx by several surrounding hydrophilic residues (Fig. 5h). Overall, the structure reveals a canonical CHD family chromatin remodeler architecture of PKL, with the ATPase domain clamping DNA for chromatin remodeling, CD1 assisting DNA binding, and CD2 connecting CD1 and ATPase domains. To confirm the structural observations, we performed *in vitro* mutagenesis studies. The mutations of the ATPase domain active site (D422N and E423Q), the DNA binding residues of the ATPase (K428E, K433E, R728E, and R426E), and the DNA binding residues of CD2 (H159D, R160E, and R164E) all significantly reduced the chromatin remodeling activity of PKL (Fig. 5i), supporting their critical roles.

### Recognition of histones by PKL

Like in most chromatin remodeler structures^58^, the H4 tail extends out from the core region with a batch of positively charged residues H4K16, H4R17, H4H18, H4R19, and H4K20 being recognized by a negatively charged PKL Lobe2 pocket formed by Glu577, Asp625, Glu628, and Asp629 through extensive hydrogen bonding and salt bridge interactions (Fig. 6a). The methyllysine of H3K4me3 is anchored by the hydrophobic residue Trp135 and Leu138 by hydrophobic and cation-π interactions in a classic mode (Fig. 6b). Since PKL recognizes H3K4me3 through its double CD, we next investigated the biochemical basis of the regulation of PKL’s function by H3K4me3. Similar to human CHD1^22^, PKL exhibits nearly identical chromatin remodeling activity towards the H3K4me3-modified and unmodified NCPs (Fig. 6c), suggesting that the H3K4me3-binding does not play a critical role in PKL’s activity. We further investigate the role of different histone tail binding sites in PKL activity. Unlike the H4-binding residue mutation (D629Q), which significantly decreases the activity of PKL, the H3K4me3 binding residue mutation of W135A possesses similar activity as the WT protein to an H3K4me3-containing NCP substrate (Fig. 6d), further strengthening that H4, but not H3, plays a critical role in the chromatin remodeling by PKL. Together with our functional data, we conclude that the H3K4me3 modification plays a role in recruiting PKL to H3K4me3-enriched target loci but does not regulate the activity of PKL.

**Fig. 6.**
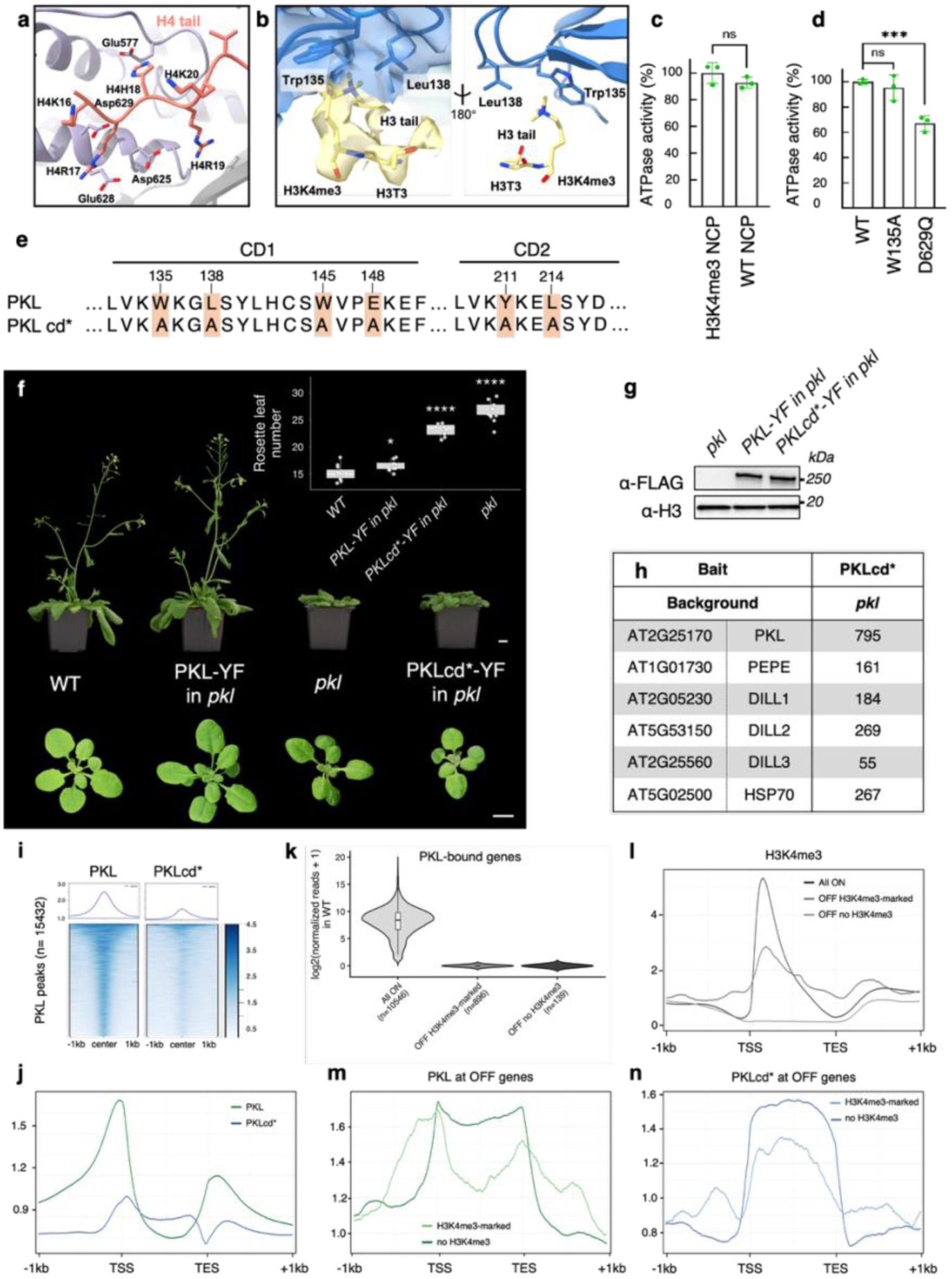
The chromodomain is recruiting PKL to H3K4me3-marked chromatin. a, Recognition of the H4 tail by PKL. **b,** The recognition of H3K4me3 by PKL with the map of H3 tail overlaid. **c,** *In vitro* activity assays of PKL towards the H3K4me3 modified or unmodified NCPs. **d,** *In vitro* assays of representative PKL mutants affecting H4 tail and H3K4me3 binding. The data are presented as means ± s.d. (*n* = 3); ***P-value < 0.001. **e,** Amino acid substitutions in the CD1 and CD2 of PKL. **f,** Plant phenotype and quantification of the juvenile-to-adult transition by counting the number of rosette leaves at the time of bolting of WT, *pkl* and *pkl* expressing YF-tagged PKL of mutated PKLcd*. Bar = 1cm. * and **** denote adjusted P-value < 0.05 and 0.0001. **g,** Western blotting of YF-tagged PKL and PKLcd*. **h,** Mean spectral counts detected by IP-MS for YF-tagged PKLcd*. **i,** Metaplot and heatmap of PKL and PKLcd* enrichment at called PKL peaks. **j,** Metaplot of PKL and PKLcd* enrichment at PKL-bound genes. **k,** Violin plot presenting the expression in WT of all PKL-bound transcribed in WT, PKL-bound repressed genes marked by H3K4me3 and PKL-bound repressed genes with no H3K4me3. **l,** Metaplot of H3K4me3 enrichment at the categories defined in k. **m,n,** Metaplot of PKL and PKLcd* enrichment at all expressed genes in WT, repressed genes marked by H3K4me3 and repressed genes with no H3K4me3.

### The chromodomains are required for PKL recruitment to H3K4me3-marked chromatin

To test the importance of H3K4me3 binding for PKL recruitment to chromatin *in vivo*, we transformed *pkl* plants with YF-tagged PKL carrying alanine mutations in conserved key residues of the double CD (W135A, L138A, W145A, E148A, Y211A and L214A) that contribute to the aromatic H3K4me3-binding pocket^12^ (Fig. 6b,e). Mutated *PKLcd\** did not complement the *pkl* phenotype, indicating that PKL functionality was compromised (Fig. 6f,g). While IP-MS revealed that PKLcd* still interacted with PEPE, DILL1-3 and HSP70 (Fig. 6h), ChIP-seq showed that the enrichment at PKL binding sites was strongly decreased (Fig. 6i). Instead of localizing to the nucleosome-free region, PKLcd* was enriched over gene bodies of PKL-bound genes, indicating that the ability to bind H3K4me3 is essential for PKL positioning (Fig. 6j). To test if the presence of H3K4me3 was sufficient to position PKL, we divided PKL-bound genes that are repressed in WT into two groups either marked or not by H3K4me3 and found that PKL but not PKLcd* enrichment shifted from gene bodies towards the nucleosome-free region (Fig. 6k-n). PEPE and DILL1-3 were also repositioned towards the promoter in the presence of H3K4me3 (Supplementary Fig. 9a-d). In summary, the double CD is essential for the recruitment of DILL-PKL-PEPE complexes to H3K4me3-marked chromatin and for PKL function.

## Discussion

Our study reveals multisubunit protein complexes centered on the CHD remodeler PKL that, through H3K4me3 binding, bridge histone methylation to chromatin remodeling. Assembly into larger regulatory complexes is a recurring feature of CHD remodelers. However, the association of DNAJ proteins and HSP70 chaperones with chromatin remodelers remains poorly documented, and our findings raise the possibility that similar chaperone-remodeler couplings exist more broadly but have gone undetected in other systems. Indeed, the HSP40 co-chaperone DNAJB6 co-enriches with the C-terminal disordered regions of most human CHD paralogs^59^, and in yeast, the HSP40 YDJ1 cooperates with chaperones and remodelers in nucleosome eviction at the *GAL* genes and, more broadly, at promoters genome-wide^60,61^. As HSP70 chaperones are generic ATPases whose client specificity is conferred by a diverse family of DNAJs^46,49,50^, DILL1-3 may function to bind and deliver PKL to HSP70, promoting PKL-mediated remodeling through protein folding, complex assembly, or stimulation of ATPase activity. Given that DILL1-3 and PKL co-localize genome-wide, HSP70 likely acts not only in the nucleoplasm but also on chromatin-bound DILL-PKL-PEPE complexes.

The assembly of DILL1-3 and PKL into three mutually exclusive complexes that bind and regulate shared targets supports functional redundancy among the DILLs. Three independent routes to the same PKL–HSP70 recruitment step would buffer the system against loss-of-function mutations, environmental stress, or expression noise in any single DILL gene - a common evolutionary rationale for retaining paralog redundancy without requiring functional divergence. Nevertheless, phenotypic and transcriptional differences among single *dill* mutants indicate incomplete compensation: *dill1*, which shows the strongest phenotype, corresponds to the most highly expressed and most abundant DILL in PKL immunoprecipitates, whereas *dill3* shows a stronger phenotype than *dill2* despite lower DILL3 expression and abundance, indicating that expression or abundance alone cannot explain the partial redundancy. Confocal imaging revealed no major differences in DILL expression patterns or subcellular localization at the tissue level examined. However, ePlant expression data indicate that PKL, PEPE and DILL1-3 transcript levels vary considerably across organs and developmental stages, raising the possibility that individual DILLs are differentially deployed in specific cell types or physiological contexts not captured by our confocal analysis. Such context-specific DILL usage could help explain why PKL has been implicated in a strikingly broad range of processes, including flowering time, lateral root formation, and abiotic stress responses^25,26,27,33,35,36,39^, despite each individual DILL contributing only partially to bulk *pkl*-like phenotypes. Distinct DILLs may predominate in distinct developmental or environmental contexts. Alternatively, despite their shared null phenotype, individual DILL-PKL-PEPE complexes may have subtly distinct genomic targeting or kinetic/regulatory properties.

RNA-seq showed that most PKL-bound genes are expressed, and that PKL targets are both up- and down-regulated in *pkl*, indicating that PKL acts as an activator or repressor in a locus-dependent manner. How does this H3K4me3-reading mechanism relate to PKL’s long-established genetic antagonism with Polycomb Repressive Complex 2 (PRC2) and its association with the repressive mark H3K27me3?^25,26,27,35^ We propose that PKL employs at least two distinct, locus-context-dependent targeting logics rather than a single unified mechanism. At loci undergoing PRC2-mediated silencing during developmental transitions, sequence-specific repressors such as VAL1 and VAL2 recruit PKL independently of histone marks^39^, where its ATPase activity locally increases nucleosome density at H3K27me3 spreading - but not nucleation - regions, generating unmodified nucleosome substrate that promotes PRC2 spreading and the stable inheritance of Polycomb silencing through cell division^23^. This mode of action depends on PKL’s SANT-SLIDE DNA-binding module and its reported association with the PRC2 subunit MSI1^23^, rather than on chromodomain-mediated H3K4me3 recognition. At actively transcribed loci, by contrast, the chromodomain–H3K4me3 mechanism described here positions DILL-PKL-PEPE complexes directly at promoters to fine-tune transcription bidirectionally, independently of PRC2. Under this model, the genetic antagonism between PKL and PRC2 reported previously^25,26^ may not reflect direct competition for the same chromatin substrate, but rather PKL’s dual role: reinforcing Polycomb memory at silenced developmental loci while independently fine-tuning transcription at active genes elsewhere in the genome.

## Methods

### Plant materials and growth conditions

All *Arabidopsis thaliana* plants were in the Col-0 background. T-DNA lines were obtained from the European Arabidopsis Stock Centre: *pkl-10* (GABI_273E06), *pepe* (SK28293), *dill2* (GABIseq_431G03) and *dill3* (SAILseq_501_C08). Plants were grown under 16 h light/8 h dark at 21°C (140 μmol m⁻² s⁻¹) after 2 days of stratification at 4°C. Plants for IP-MS, phenotyping and flowering-time measurements were grown on soil. Seedlings for immunoblotting, RNA-seq and ChIP-seq were grown on 1% agar plates (½ MS) for 10 days after a 2-day germination period. Seedlings for microscopy were grown vertically for 5-10 days.

### Flowering time measurements

Rosette leaves were counted for 20 plants per genotype after 4 weeks of soil growth. Genotypes were compared to WT using Welch’s two-sample *t*-test, with Benjamini-Hochberg correction for multiple testing; adjusted P < 0.05, 0.01 and 0.001 are indicated by one, two and three asterisks, respectively.

### Generation of translational fusion lines

PKL-YF (*pPKL::gPKL::2xYFP-3xFLAG::3’utrPKL*) was generated by bacterial recombineering into a transformation-competent artificial chromosome^40,41^ and introduced into *pkl-10* by floral dip^63^. PEPE-, DILL1-, DILL2- and DILL3-YF constructs were assembled by Gibson and Gateway cloning, combining an entry backbone (pENTRY22113), the PCR-amplified 5’UTR/promoter and coding sequence (minus stop codon) of each gene, a YF tag with stop codon, and the 3’UTR, then transferred into pGWB501 (Addgene) by LR reaction (Thermo Fisher) and transformed into the corresponding mutant background. All mutations were introduced by PCR. T1 plants were selected on hygromycin and genotyped by qPCR (90 plants per construct) using YF tag-specific primers. Single-insertion lines were confirmed by PCR across the YF-gene junction and by western blot of rosette leaf extracts. Primers are listed in Supplementary Data 6.

### CRISPR/Cas9 cloning

The *dill1* mutant was generated by introducing two sgRNA protospacers into pAGM55261, which expresses an intronized maize Cas9 (zCas9i)^64^. A construct containing the protospacers and flanking sgRNA regulatory elements was PCR-amplified from pHEE2E-TRI using protospacer-specific primers (Supplementary Data 6) and cloned into pAGM55261 by BsaI-based Golden Gate assembly. Protospacer sequences were selected using CHOPCHOP^65^ or designed manually.

### Phylogenetic analysis and protein alignments

PEPE homologs were identified by BLASTP (NCBI Protein, Phytozome; identity ≥30%, coverage ≥50%, E-value ≤1e−5) across representative monocot, dicot and basal land-plant species: *Arabidopsis lyrata* (AL1G10610.t1), *Brassica rapa* (I05633.1.p), *Vitis vinifera* (VIT_215s0046g02890.1), *Solanum lycopersicum* (Solyc05g049880.3.1), *Populus trichocarpa* (Potri.014G082600.1.p), *Oryza sativa* (LOC_Os02g44540.3), *Brachypodium distachyon* (Bradi3g51040.1.p) and *Zea mays* (Zm00001d017523_P002). Sequences were aligned in ClustalW^66^ (default gap penalties, BLOSUM matrix), and a maximum-likelihood tree was built in MEGA X^67^ (100 bootstrap replicates), rooted on *Marchantia polymorpha*. DUF3444-containing J-domain proteins were identified using InterPro.

### Immunoprecipitation followed by mass spectrometry

IP-MS was performed in duplicate per genotype. Proteins were extracted from 5 g flower buds, and nucleic acids digested with benzonase (Sigma; 0.5 μL/100 mg tissue). FLAG-tagged proteins were immunoprecipitated with FLAG M2 antibody (Sigma F1804) and protein A/G magnetic beads (Thermo), then eluted, precipitated and digested with 500 ng sequencing-grade trypsin in 20 mM Tris, 2 mM CaCl2, 2 mM TCEP, 15 mM chloroacetamide, pH 8.2 overnight at 37°C. Samples were analyzed by LC-MS/MS on a timsTOF Pro (Bruker) coupled to an Evosep One, using the 15 samples/day method (PSC-15-100-3-UHPnC column, 50°C). MS spectra were acquired in ddaPASEF mode (m/z 100–1700; ion mobility 1/K0 0.60–1.60 Vs/cm²; 166 ms accumulation/ramp time; 1 survey scan followed by 10 PASEF ramps; 1.9 s cycle time), excluding singly charged ions, with MS/MS isolation windows of m/z 2.0 (<700) and 3.0 (>700). Data were processed in FragPipe v17 (MSFragger) against the *Arabidopsis thaliana* TAIR10 database, with oxidation (M) as variable and carbamidomethylation (C) as fixed modifications. Proteins with ≥2 peptides and nonzero MS1 intensity were considered true hits.

Differential enrichment was assessed for proteins detected in either PKL or WT (n = 529) using a two-sided Student’s *t*-test on log2-transformed spectral counts (n = 2 replicates/group); |log2FC| > 1 and P < 0.05 defined significant enrichment. HSP70-1 (AT5G02500) enrichment relative to WT was expressed as fold-change and z-score against the log2-transformed background distribution of all detected proteins (count > 0) across the 24 experiments.

### Immunoblot analysis

Proteins extracted in 40 mM Tris-HCl pH 7.5, 10% glycerol, 5 mM MgCl2, 4% SDS were quantified (BCA kit, Pierce). 10 μg was separated by SDS-PAGE, transferred, and probed with HRP-conjugated FLAG antibody (Sigma-A8592), then stripped and reprobed with H3 antibody (ab1791) and HRP-conjugated anti-rabbit IgG (Amersham NA934).

### Microscopy

Images were acquired on a Leica TCS SP8 microscope (HC PL APO 63x/1.20W motCORR CS2 objective). For leaf and floral tissues (sepals, petals, pollen, stigmas, ovules), YF constructs were excited at 514 nm (argon laser) with emission collected at 525–550 nm (descanned HyD) and transmission on a PMT-Trans channel. For roots, YF and propidium iodide were sequentially excited at 960 nm and 1040 nm (InSight DS+ Dual multiphoton lasers) with emission collected on non-descanned HyD-RLD detectors (460/50 & 525/50 with 495 SP for YF; 585/40 & 650/50 with 620 SP for propidium iodide).

### RT-qPCR and RNA-sequencing

RNA (RNeasy Plant Mini Kit, Qiagen) was reverse-transcribed (1 μg) using the iScript cDNA synthesis kit (Bio-Rad). RT-qPCR was performed on 5 μL cDNA with KAPA SYBR® FAST mastermix (Merck); melt curves were checked in Bio-Rad CFX Maestro, and relative expression was calculated as 2^−ΔΔCt^. Libraries for sequencing were prepared with the TruSeq RNA Library Prep Kit v2 and sequenced on a NovaSeq X platform (150-bp paired-end). Reads were mapped to TAIR10 (Ensembl Plants release 57, 2023-09-06) with STAR^68^ and counted with featureCounts^69^ (default parameters). Differential expression was assessed with DESeq2 (FC > 2, padj < 0.05) across four replicates per genotype. Primers are listed in Supplementary Data 6.

### Chromatin immunoprecipitation followed by sequencing

4 g of 10-day-old seedlings were crosslinked in 1% formaldehyde/1x PBS. Nuclei were resuspended in 500 mM Tris pH 8.0, 10 mM EDTA, 1% SDS, 5 mM benzamidine, 1x PIC, 1 mM PMSF, and chromatin was sheared (Bioruptor® Plus), precleared with protein A/G beads (Thermo), and immunoprecipitated with anti-GFP antibody (Thermo A-6455) and protein A/G beads. After washing, elution and crosslink reversal, RNA was removed (RNase, Thermo) and protein digested (Proteinase K, Invitrogen); DNA was purified by phenol/chloroform extraction and libraries prepared with the Ovation® Ultralow System V2 (Tecan). Sequencing was performed on a NovaSeq 6000 (100-bp single-end reads). Reads were aligned to the *Arabidopsis thaliana* TAIR10 genome with Bowtie2^70^ and peaks called against input with MACS2^71^. Peaks were visualized in IGV^72^. Two replicates were analyzed per condition.

### Protein expression and purification

PKL was cloned into a modified pFastHTB vector with an N-terminal Strep-His6 tag and expressed in Sf9 cells (Bac-to-Bac baculovirus system, Invitrogen). Recombinant protein was purified by Strep-Tactin affinity (Qiagen), followed by Heparin and Superose 6 columns (GE Healthcare); all PKL variants were purified identically. The double CD (residues 96–241) were subcloned into a modified pSumo vector (N-terminal Sumo-His6 tag), expressed in *E. coli* Rosetta (DE3) in LB medium (37°C; induced at OD^600^ = 0.8 with 0.2 mM IPTG, 18°C), and purified by Ni^2+^ affinity (GenScript), Ulp1 cleavage, a second Ni^2+^ affinity step, and a Superdex G75 column. The PHD finger (residues 48–99) was purified identically.

*Arabidopsis* histone proteins for NCP reconstitution were expressed and purified as for *X. laevis* histones^55^. H3K4me3-modified H3 was purchased (Kesheng Jingtai); the 226-bp Widom 601-derived DNA was PCR-amplified^56,73^, and NCPs were assembled by salt-gradient dialysis^73^. Peptides were synthesized by GL Biochem (Shanghai). Oligonucleotide sequences are listed in Supplementary Data 6.

### Bio-layer interferometry–based *in vitro* binding assay

Purified PKL domains were biotinylated (EZ-Link NHS-PEG4-Biotin kit, Thermo Fisher Scientific) and immobilized on Streptavidin biosensors (Sartorius). Binding was measured on an Octet RED96 (Sartorius) and analyzed using Octet Data Analysis 10.0.

### *In vitro* chromatin remodeling assay

Chromatin remodeling activity was measured via ADP production using the ADP-Glo™ Kinase Assay kit (Promega). WT or variant PKL (3.2 μM) and NCP (0.4 μM) were combined in 50 mM NaCl, 5 mM MgCl2, 1 mM ATP, 25 mM Tris-HCl pH 7.5; after 60 min, luminescence was read on a Synergy H1 plate reader (BioTek). Assays were performed in triplicate and analyzed in GraphPad Prism 9.

### Cryo-EM grid preparation and data collection

PKL (9 μM) and NCP (1.8 μM) were mixed with 0.5 mM ADP, 16 mM NaF and 2 mM BeSO4, then stabilized by GraFix using top (25 mM NaCl, 1 mM MgCl2, 10% glycerol, 0.01% glutaraldehyde, 10 mM HEPES pH 7.4) and bottom (25 mM NaCl, 1 mM MgCl2, 30% glycerol, 0.15% glutaraldehyde, 10 mM HEPES pH 7.4) solutions, both supplemented with 0.5 mM ADP, 0.8 mM NaF and 1 mM BeSO4. Samples were ultracentrifuged (35,000 rpm, 18 h, 4°C; Beckman SW-41Ti), and the migrated peak fraction was dialyzed into 25 mM NaCl, 2 mM MgCl2, 3 mM DTT, 1 mM ADP, 16 mM NaF, 2 mM BeSO4, 20 mM Tris-HCl pH 7.5, then concentrated to 3 mg/mL.

Quantifoil Cu R1.2/1.3 300-mesh grids (Micro Tools GmbH) were glow-discharged (45 s, PDC-32G, Harrick Plasma), loaded with 4 μL sample, blotted (2 s) and vitrified in liquid ethane (Vitrobot Mark IV, FEI; 4°C, 100% humidity). Data were collected on a Titan Krios (FEI/Thermo, 300 keV; SUSTech Cryo-EM Center) with a K3 detector and GIF Quantum energy filter, using EPU at 105,000× nominal magnification in super-resolution mode (0.855 Å pixel size). Micrographs were dose-fractionated into 32 frames (24 counts/physical pixel/s; total dose ∼50 e⁻/Å²).

### Cryo-EM data processing and structure determination

Data were processed in CryoSPARC. 8,643 movies were split into two subsets for patch motion correction and CTF estimation. Particles were picked using templates from the Chd1-nucleosome structure (EMD-51247)^57^, yielding 2,763,846 and 2,834,398 particles per subset; 2D classification removed low-quality particles (646,499 and 769,574 remaining). Ab initio reconstruction from one subset generated templates for heterogeneous refinement of the combined particle set, which separated into three classes, including the 1:1 and 2:1 PKL-NCP complexes. The 1:1 class was refined by non-uniform then local refinement (3.14 Å); the 2:1 class underwent an additional 2D classification round before non-uniform refinement (3.05 Å). The AlphaFold-derived model was fit into the map and manually rebuilt in Coot^44,74^, with real-space refinement in Phenix using Ramachandran restraints. Structure figures were generated in UCSF ChimeraX^75^.

## Data availability

The structures have been deposited in the Protein Data Bank with the accession codes 27CG and 27CF. The cryo-EM maps have been deposited in the Electron Microscopy Data Bank with accession codes EMD-81049 and EMD-81050. RNA-seq and ChIP-seq datasets were deposited into ArrayExpress with the accession number E-MTAB-17563 and E-MTAB-17537 respectively. IP-MS data were deposited in the PRIDE database with the dataset identifier PXD083022.

## Acknowledgment

We thank the staff members at the Functional Genomics Center Zürich and at the SUSTech Cryo-EM Center for their assistance with data collection. This work was supported by the Guangdong Major Project of Basic Research (2026B0303000001 to J.D.), National Natural Science Foundation of China (32325008 to J.D.), Shenzhen Science and Technology Program (RCJC20221008092720004 to J.D.), China Postdoctoral Science Foundation (BX20250129 to G.X.), the Swiss National Science Foundation (PP00P3_1176957, PP00P3_176957, and PP00P3_217600 to S.B.), the European Union’s Horizon 2020 research and innovation program (847585_ESR14 to S.B.), the Novartis Foundation for medical-biological Research (23A068 to S.B.), the Forschungskredit der Universität Zürich (FK-21-134 to E.R.) and the Olga Mayenfisch Stiftung (Forschungsprojekt to S.B.). J.D. is an investigator of the SUSTech Institute for Biological Electron Microscopy.

## Author contributions

S.B. and J.D. conceived the study and designed the experiments. S.C., A.L., E.R., N.C.H., C.C., A.S., E.P., A.G.F., S.P., L.C., D.K., G.X., R.L., J.A.L and S.B. performed the experiments, collected and analyzed the data. All authors contributed to data interpretation and reviewed the manuscript.

## Declaration of generative AI use

The authors used Claude for language editing and polishing, after which the authors carefully reviewed and edited the content and take full responsibility for the published article.

