## Supplementary Figures for "H3K4me3 recruits the chromatin remodeling DILL-PICKLE-PEPPERCORN complexes to promoters in plants"

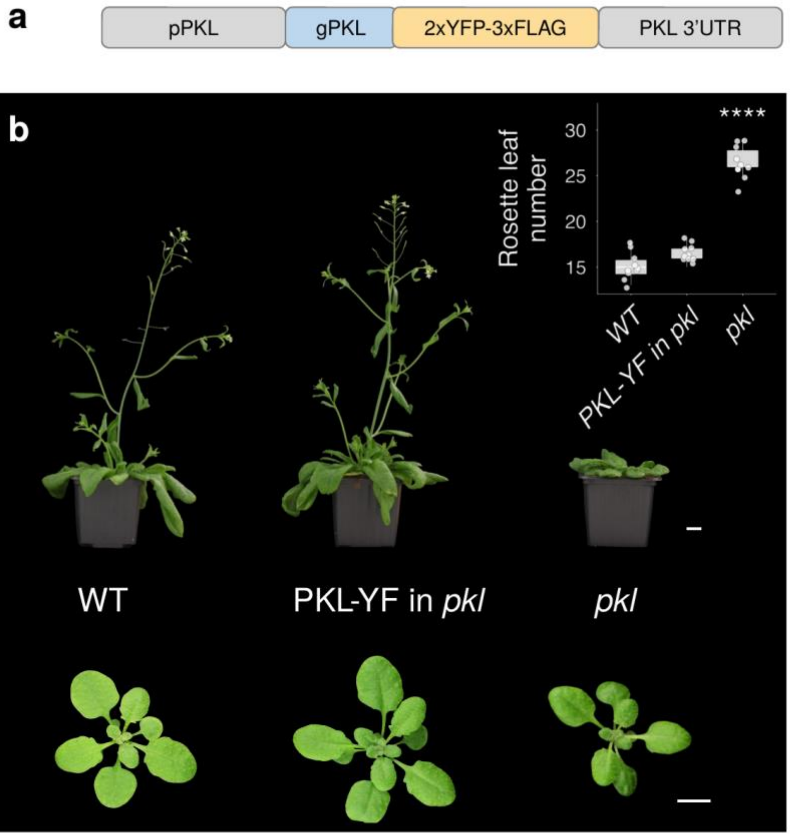


**Supplementary Figure 1. Complementation of the *pkl-10* mutant. a,** Construct schema of 2xYFP-3xFLAG (YF) tagged genomic PKL driven by its endogenous promoter. **b,** Plant phenotype and quantification of the juvenile-to-adult transition by counting the number of rosette leaves at the time of bolting of WT, *pkl* and complemented *pkl* mutant expressing YF-tagged PKL. Bar = 1cm. **** denote adjusted P-value $< 0.0001$.

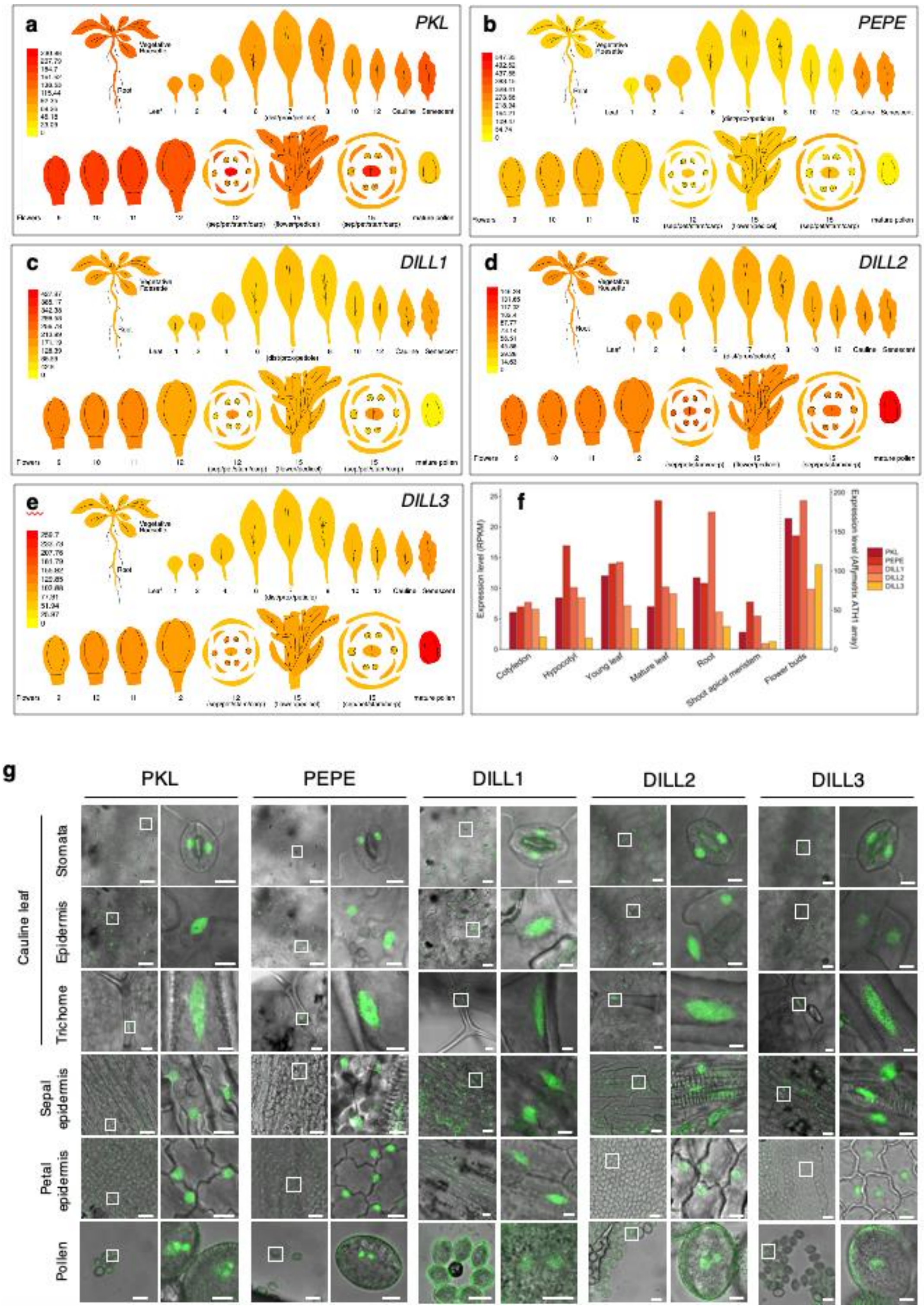


**Supplementary Figure 2. PKL is ubiquitously expressed. a-e,** Relative expression levels of the *PKL*, *PEPE* and *DILL1-3* genes in select tissues from ePlant expression viewers. **f,** Gene expression of *PKL*, *PEPE* and *DILL1-3* in cotyledons, hypocotyls, young and mature leaves, roots, shoot apical meristems and flower buds. Data downloaded from ePlant expression database. **g,** Confocal microscopy showing the localization of 2xYFP-3xFLAG (YF) tagged proteins driven by their respective endogenous promoter in various tissues. Bar = 5 µm.

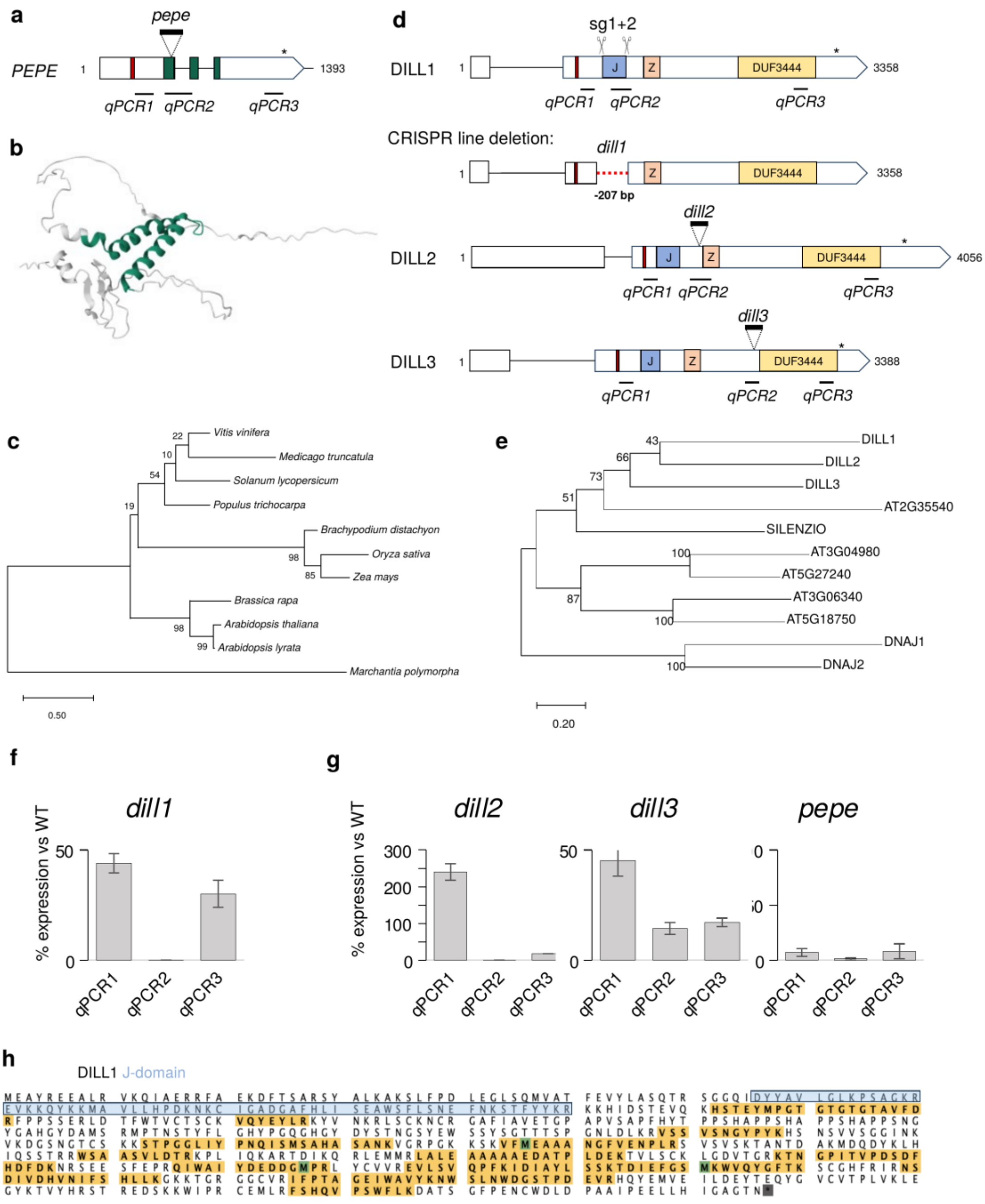


**Supplementary Figure 3. Characterization of *pepe* and *dill1-3* mutant lines. a,d** Exon/intron models of the *PEPE* and *DILL1-3* genes showing the location of T-DNA insertion, sgRNAs, the obtained CRISPR deletion and sequences amplified by RT-qPCR. J-domain (J), Zinc finger-like region (Z) and DUF3444 are highlighted. **b,** AlphaFold structure prediction of PEPE. The helix-turn-helix is in green. **c,** Phylogenetic tree of *PEPE* homologues across plant species. **e,** Protein tree of *Arabidopsis* J-domain proteins carrying a DUF3444. SILENZIO, DNAJ1 and DNAJ2 serve as references for J-proteins involved in epigenetic gene regulation. **f,g,** RT-qPCR quantifying the expression of *PEPE*, *DILL1-3* in their respective mutant background versus WT. **h,** DILL1 peptides identified by IP-MS (in yellow) with PKL-YF in *dill1*. The J-domain is highlighted in blue.

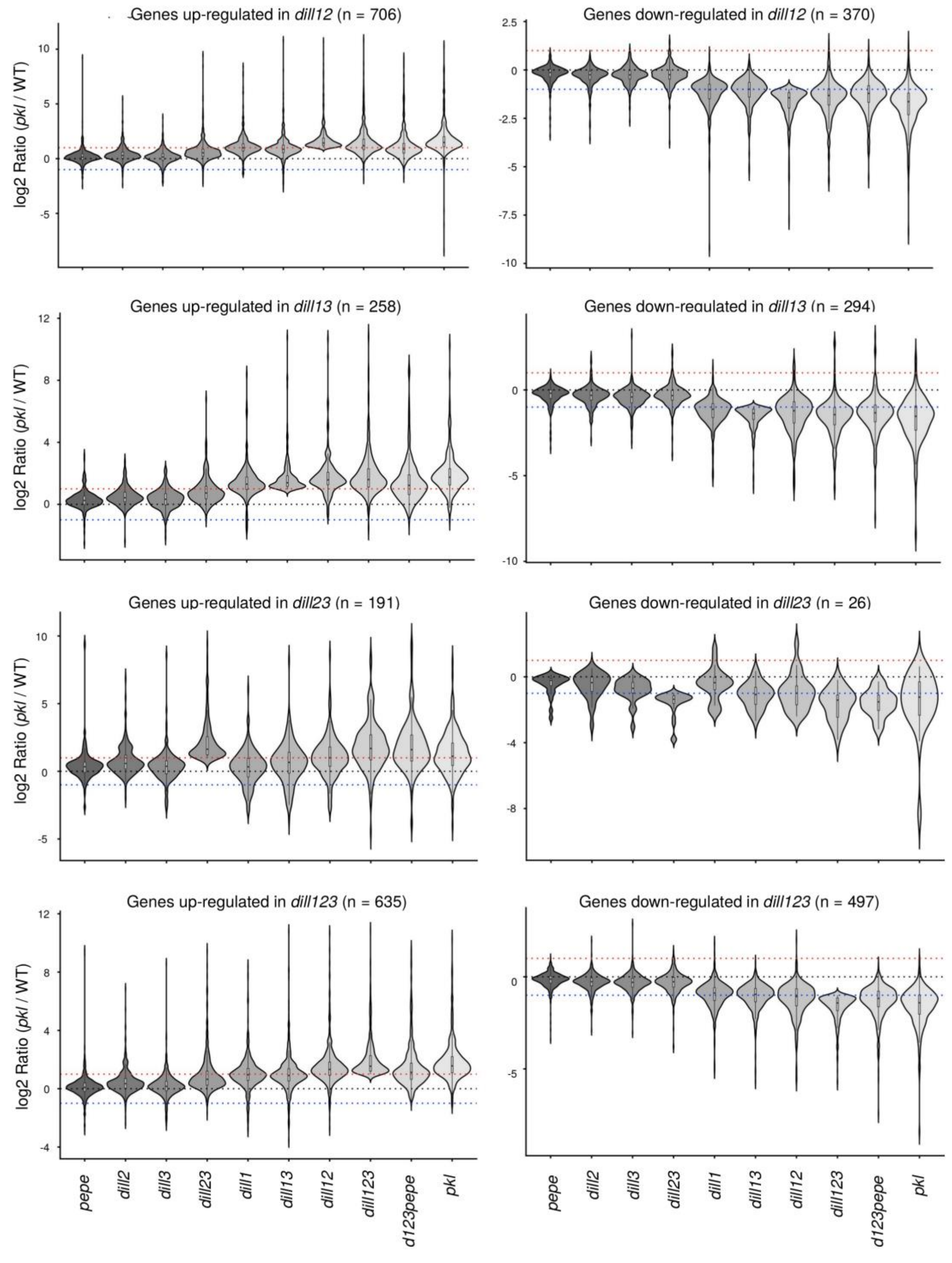


**Supplementary Figure 4. DILL1-3 are functionally redundant with PKL.** Violin plots showing gene expression in *pepe*, *dill1-3, pkl* and high-order mutant combinations for the genes differentially regulated in *dill12*, *dill13*, *dill23* and *dill123* (fold-change > 2 and P-value < 0.05).

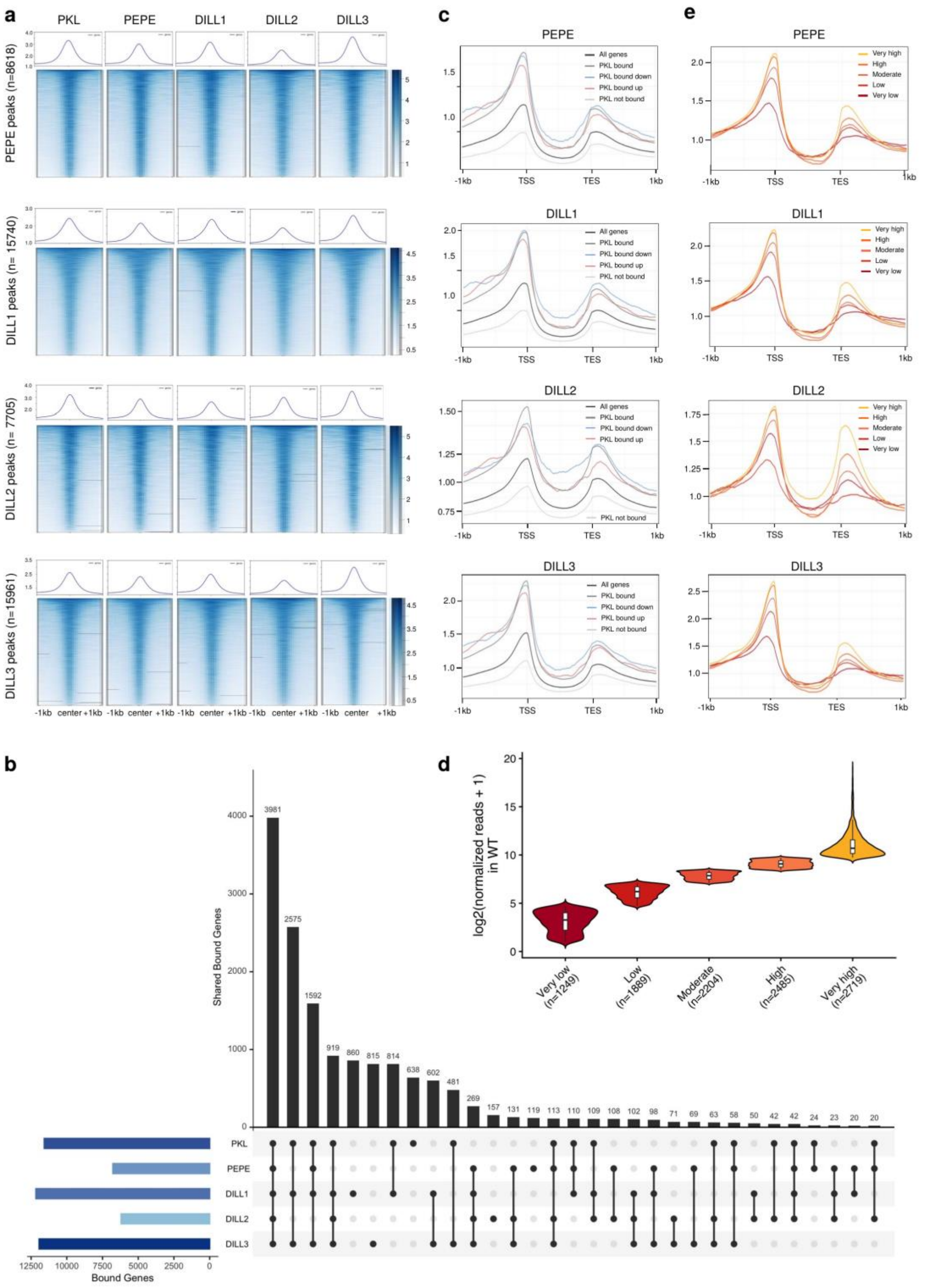

**Supplementary Figure 5. PEPE and DILL1-3 co-localize at active promoters. a,** Metaplot and heatmap of PKL, PEPE and DILL1-3 at PEPE and DILL1-3 called peaks. **b,** Upset plot showing the overlap between genes bound by PKL, PEPE and DILL1-3. **c,** Metaplot of PEPE and DILL1-3 at all protein-coding genes, PKL-bound, PKL–unbound and, up- and down-regulated genes in *pkl*. **d,** Violin plot presenting the expression of all actively transcribed genes in WT bound by PKL binned into five categories. **e,** Metaplot of PEPE and DILL1-3 enrichment in the five groups of active genes in WT defined according to their expression level.

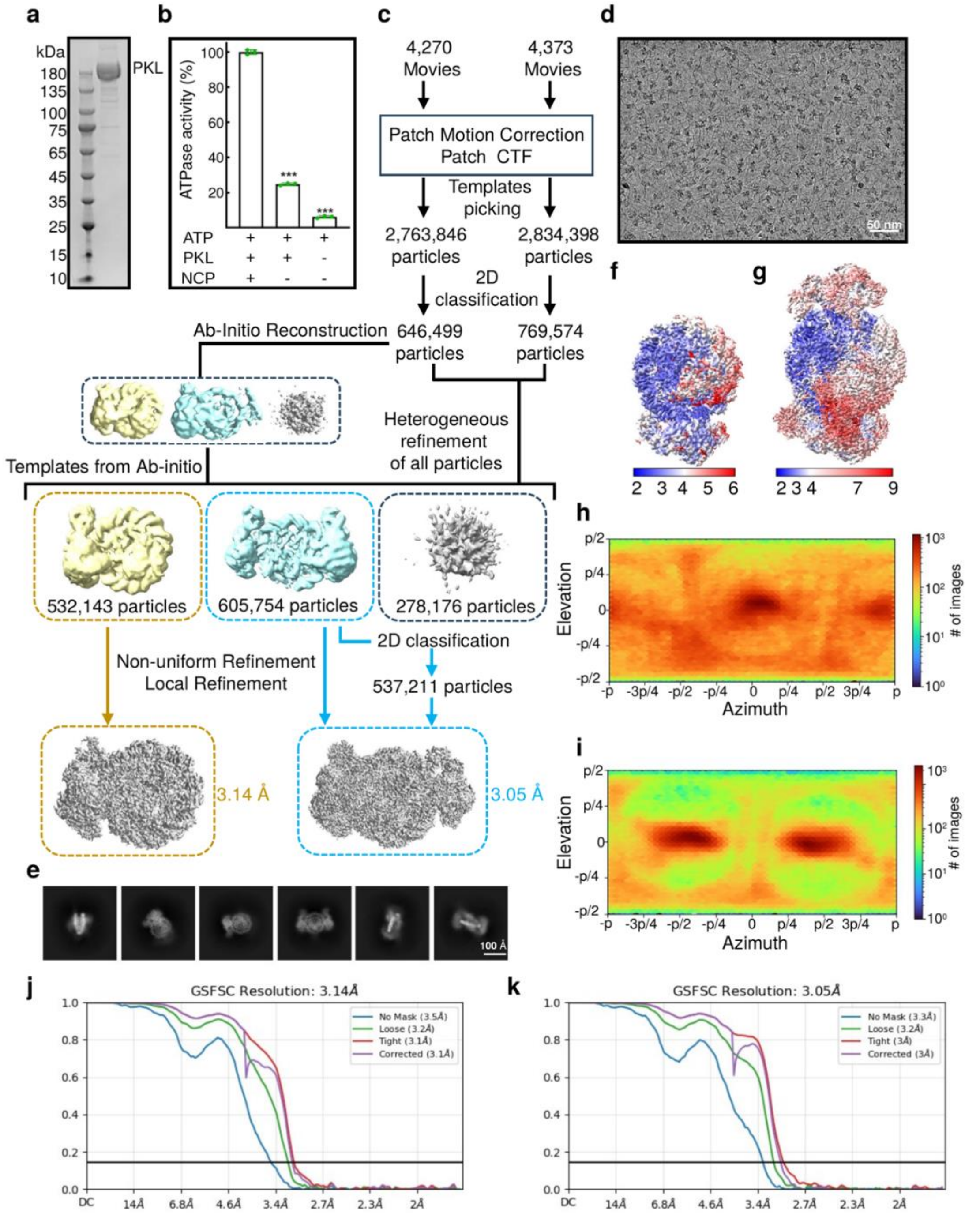


**Supplementary Figure 6. Cryo-EM structural analysis of PKL-NCP complex. a,** The SDS-PAGE of purified PKL. **b,** The ATPase activity of PKL, data are mean ± SD (n = 3 independent experiments); ***P < 0.001 (two-tailed Student's t-test). **c,** Flow chart of cryo-EM data processing. **d,** Representative micrograph of cryo-EM data collection with scale bar (50 nm). **e,** Representative 2D classes of PKL-NCP complex. **f-g,** Local resolution map of the 1:1 complex (f) and 2:1 complex (g). **h-i,** Angular distribution of particles for the final 3D map of the 1:1 complex (h) and 2:1 complex (i). **j-k,** FSC curves of the 1:1 complex (j) and the 2:1 complex (k).

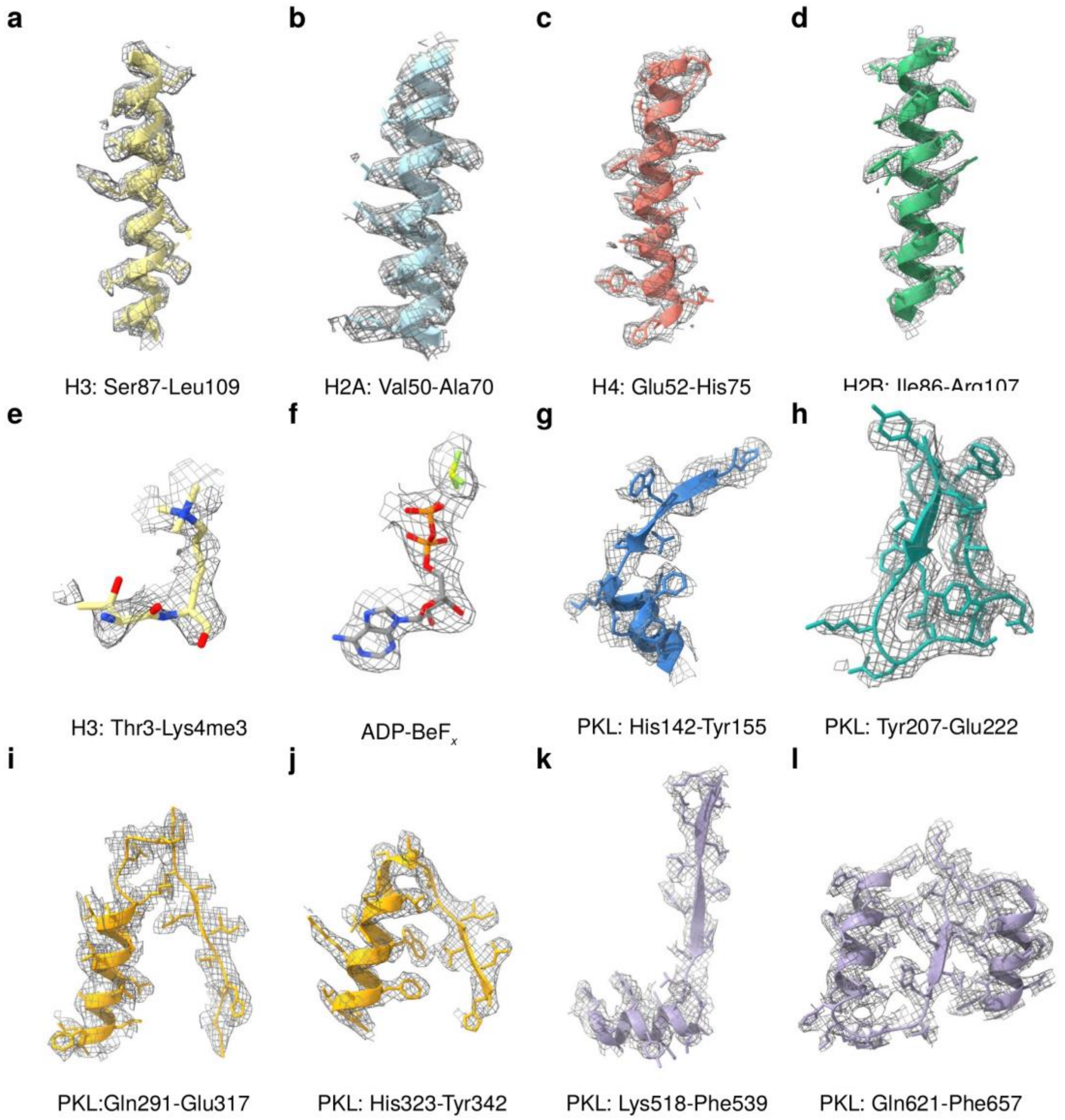


**Supplementary Figure 7. Cryo-EM map fitting for representative regions of PKL-NCP complex in ADP-BeFx bound form. a-l,** Electron density maps showed the fitting of histones (a-e), ADP-BeF*x* (f), and representative PKL regions (g-l).

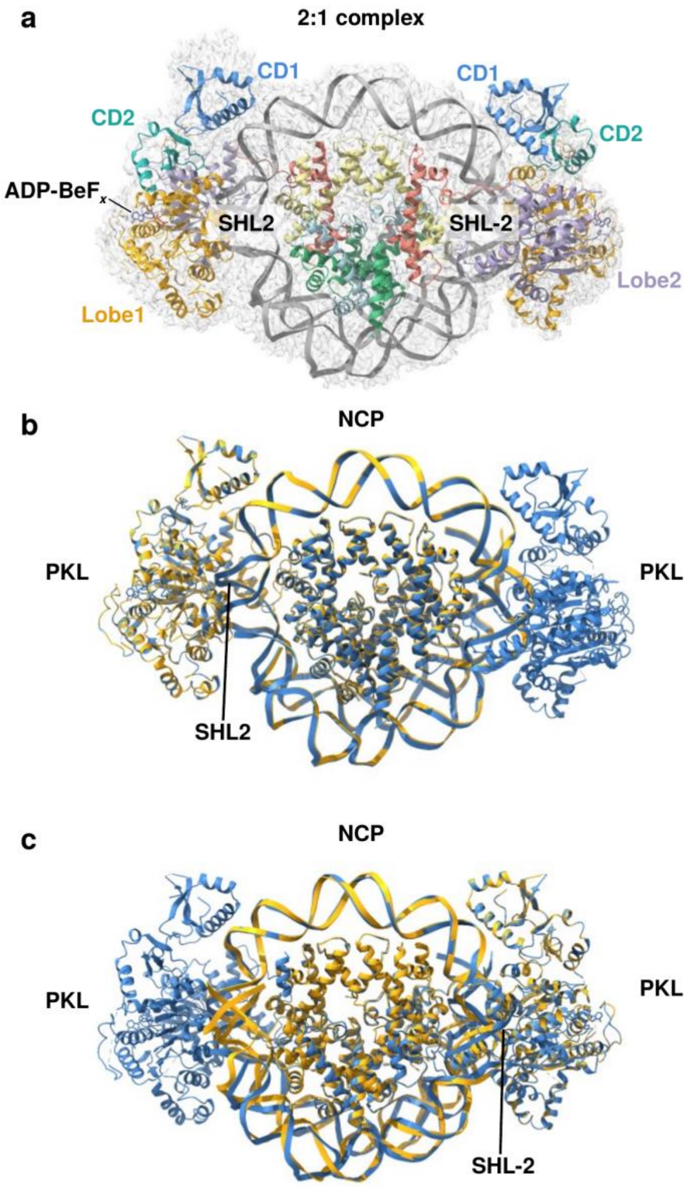


**Supplementary Figure 8. Cryo-EM structure of 2:1 complex of PKL-NCP in ADP-BeFx bound state. a,** The structure of the 2:1 complex PKL-NCP in ADP-BeF*x*-bound state with PKL bound to SHL2 and SHL-2 position of NCP. The cryo-EM map is overlaid. **b-c,** Superimpositions of the 1:1 complex (yellow) with the PKL at SHL2 (b) and SHL-2 (c) positions in the 2:1 complex reveal similar overall structures and NCP-binding modes.

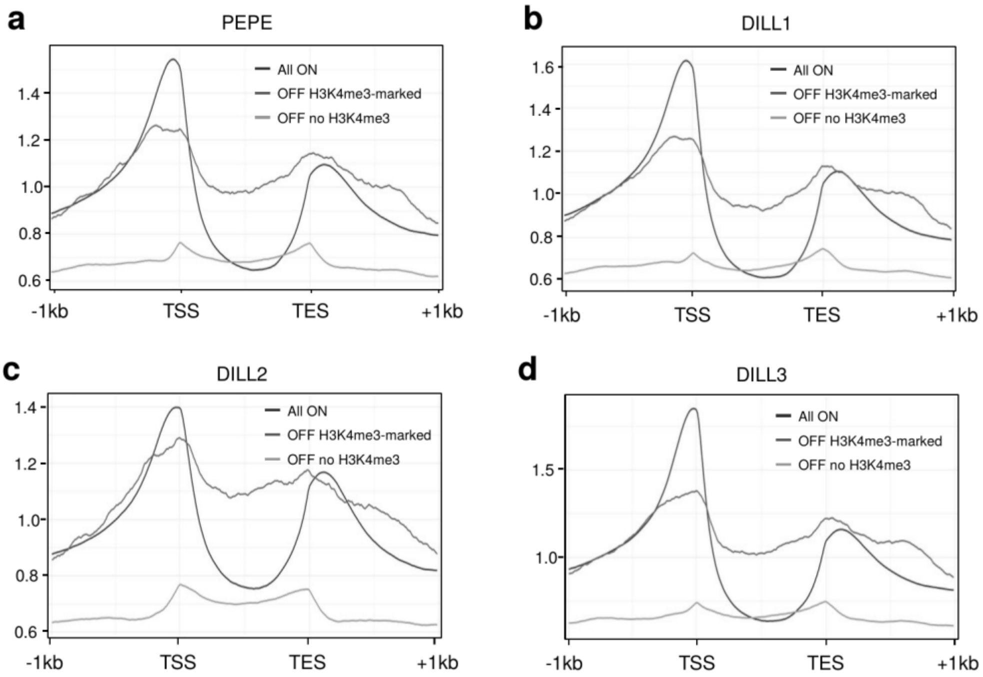


**Supplementary Figure 9. H3K4me3 positions PEPE and DILL1-3. a-d,** Metaplot of PEPE and DILL1-3 enrichment at all PKL-bound genes expressed in WT, PKL-bound repressed genes marked by H3K4me3 and PKL-bound repressed genes with no H3K4me3.
